# Viral mediated knockdown of GATA6 in SMA iPSC-derived astrocytes prevents motor neuron loss and microglial activation

**DOI:** 10.1101/2021.10.11.463972

**Authors:** Reilly L. Allison, Emily Welby, Guzal Khayrullina, Barrington G. Burnett, Allison D. Ebert

## Abstract

Spinal muscular atrophy (SMA), a pediatric genetic disorder, is characterized by the profound loss of spinal cord motor neurons and subsequent muscle atrophy and death. Although the mechanisms underlying motor neuron loss are not entirely clear, data from our work and others support the idea that glial cells contribute to disease pathology. GATA6, a transcription factor that we have previously shown to be upregulated in SMA astrocytes, is negatively regulated by SMN and can increase the expression of inflammatory regulator NFκB. In this study, we identified upregulated GATA6 as a contributor to increased activation, pro-inflammatory ligand production, and neurotoxicity in spinal-cord patterned astrocytes differentiated from SMA patient induced pluripotent stem cells. Reducing GATA6 expression in SMA astrocytes via lentiviral infection ameliorated these effects to healthy control levels. Additionally, we found that SMA astrocytes contribute to SMA microglial phagocytosis, which was again decreased by lentiviral-mediated knockdown of GATA6. Together these data identify a role of GATA6 in SMA astrocyte pathology and further highlight glia as important targets of therapeutic intervention in SMA.

## Introduction

Spinal muscular atrophy (SMA) is a prevalent genetic cause of infant mortality with an incidence of 1 in every 6,000-10,000 births.^1,2^ SMA is caused by a loss of function mutation in the survival motor neuron-1 (*SMN1*) gene, which leads to reduced SMN protein expression.^3^ This leads to the selective loss of motor neurons in the spinal cord and the atrophy of skeletal muscle, ultimately resulting in respiratory distress and death.^4^ A second gene, *SMN2*, produces some functional SMN, but due to alternative splicing this gene mainly produces a truncated, non-functional protein (SMNΔ7). SMN has been shown to be ubiquitously expressed in all cell and tissue types, though its loss in SMA particularly impacts motor neurons.^5^ SMA motor neurons (MNs) show intrinsic deficits in splicing and electrophysiological function, however restoring SMN to motor neurons alone does not significantly improve disease pathogenesis.^6–9^ Currently, there are 3 FDA approved treatments for SMA, two of which focus on correcting or adjusting the splicing of *SMN2* (nusinersen, risdiplam), as well as a gene therapy to deliver functional *SMN1* (onasemnogene abeparvovec).^10^ With recent studies finding potential risk in the AAV9-SMN gene therapy treatment for less severe SMA patients, the clinical trials to expand this treatment have halted.^11–13^ The need to identify additional drivers of MN loss remains essential for SMA patient longevity.

Glial cells—including astrocytes and microglia—also lose SMN expression in SMA, which results in aberrant activation. Astrocytes are the most abundant cell in the central nervous system (CNS), and microglia are its primary immune cell. In normal physiological function, both have neuroprotective effects and encourage neuronal development, differentiation, and function. Specifically, astrocytes produce and release growth factors like nerve growth factor (NGF), brain derived neurotrophic factor (BDNF), and glial derived neurotrophic factor (GDNF), maintain extracellular concentrations of ions (H+, K+), free radicals, and neurotransmitters, and provide structural support.^14,15^ Microglia regulate pro- and anti-inflammatory processes and respond to pathological insults through phagocytosis, but also secrete BDNF, provide metabolic intermediates and neurotransmitters, and respond to neuronal firing through the retraction and extension of their highly motile processes.^16–18^ The emerging idea of a quad-partite synapse emphasizes the tightly interdependent and highly responsive nature of communication via secreted factors between astrocytes, microglia, and neurons in the CNS.^19^

It is well known that astrocytes and microglia activate in response to stress or injury, and upregulate pro-inflammatory cytokines, downregulate neurotrophic factor secretion, and can induce neuron death.^20^ In disease conditions like SMA, however, the changed interactions between these cells and their contribution to neuronal loss have not yet been fully characterized. Importantly, astrocyte secreted factors that are known to impact motor neuron health and development, such as glial fibrillary acidic protein (GFAP) and pro-inflammatory cytokines interleukin 1β (IL1B) and interleukin 6 (IL6), are also received by microglial membrane receptors.^21,22^ The activity of these canonical receptors, including IL1R1 and IL6R, can serve to start inflammatory mechanisms and transform ramified microglia with small cell bodies and extensive, ramified branches into large, ameboid cells capable of phagocytosis.^23–25^ Complement factors serve as “eat me” tags to activated microglia, and complement proteins C1q and C3 have been found to directly contribute to the loss of viable neurons through primary phagocytosis (phagoptosis) in several neurodegenerative diseases.^26–29^ In this way, secreted factors which are essential for astrocytes and microglia to communicate in normal physiological conditions may serve as pathological drivers of activation when dysregulated.

Our lab has previously shown that astrocytes in SMA show activation before overt motor neuron loss and can induce detrimental effects on motor neuron cell health and further astrocyte activation.^30,31^ One potential explanation for this effect is the noted change in astrocyte secreted factors, as SMA astrocytes show reduced GDNF release, abnormal microRNA production, and increased NFκB expression.^30–34^ GATA6, a zinc finger transcription factor negatively regulated by SMN and associated with NFkB,^35,36^ was accordingly found to be increased in SMA mouse and human samples.^36^ Microglial activation has been noted late in the pathology of SMA mouse models,^37^ but recent evidence demonstrates microglial phagocytosis of complement-tagged synapses on proprioceptive neurons in the SMA mouse spinal cord, indicating a role during disease progression.^38^ It is not yet known how upregulated GATA6 changes the astrocyte phenotype in SMA, nor how this impacts astrocyte interactions with motor neurons and microglia. In this study, we utilized spinal-cord patterned astrocytes differentiated from SMA patient and healthy control induced pluripotent stem cells (iPSCs) to investigate the astrocytic role of GATA6 in aberrant activation and neurotoxicity as well as astrocyte-driven microglia activation *in vitro*. We observed an increase in pro-inflammatory factor production in astrocytes associated with increased GATA6 expression. Lentiviral mediated GATA6 overexpression was found to increase motor neuron cell loss in vulnerable SMA cultures, whereas lentiviral mediated knockdown ameliorated this effect. SMA astrocytes were also found to increase activation and phagocytic ability of SMA microglia, and again GATA6 astrocyte knockdown ameliorated these effects. Together these data indicate a role for GATA6 as a major contributor to the aberrant activation and pathological functions of glial cells in SMA.

## Results

### Differentiation of spinal cord astrocytes from induced pluripotent stem cells

Though astrocytes in general share a common goal of supporting neuronal function, subtle differences in gene expression, morphology, and function in astrocytes between regions of the brain and spinal cord have been identified.^39,40^ To best model astrocyte subtypes that would interact most directly with degenerating motor neurons in SMA, we differentiated ventral-caudal patterned neural progenitor cells (NPCs) into spinal cord-like astrocytes. We utilized several recently updated astrocyte protocols to optimize our own culture conditions.^41–44^

Human induced pluripotent stem cells (iPSCs) from SMA patients and healthy (unaffected) individuals, hereafter referred to as “SMA” and “Ctrl,” were differentiated into neural progenitor cells (NPCs) through the addition of SMAD inhibitors (SMADi) with retinoic acid (RA) and smoothened agonist (SAG) to mimic spinal cord development (Figure 1A). Plating NPCs at a high density was necessary to achieve robust astrocyte differentiation (Figure 1B; NPC P2 Day 17). These NPCs were found to express known NPC-associated genes such as NESTIN, SOX2, and PAX6 (Figure 1C). They also expressed spinal cord ventral-caudal associated genes including ISL1, NKX6.1, OLIG2, HOXB4, and HOXA4, which showed similar gene expression trends to post-mortem spinal cord from SMA patients and unaffected individuals (Figure 1C). These spinal cord genes were not upregulated in SMADi only treated NPCs, which instead showed increased expression of forebrain associated genes (TBR2, SIX3, OTX2, Figure 1C).

**Figure 1.**
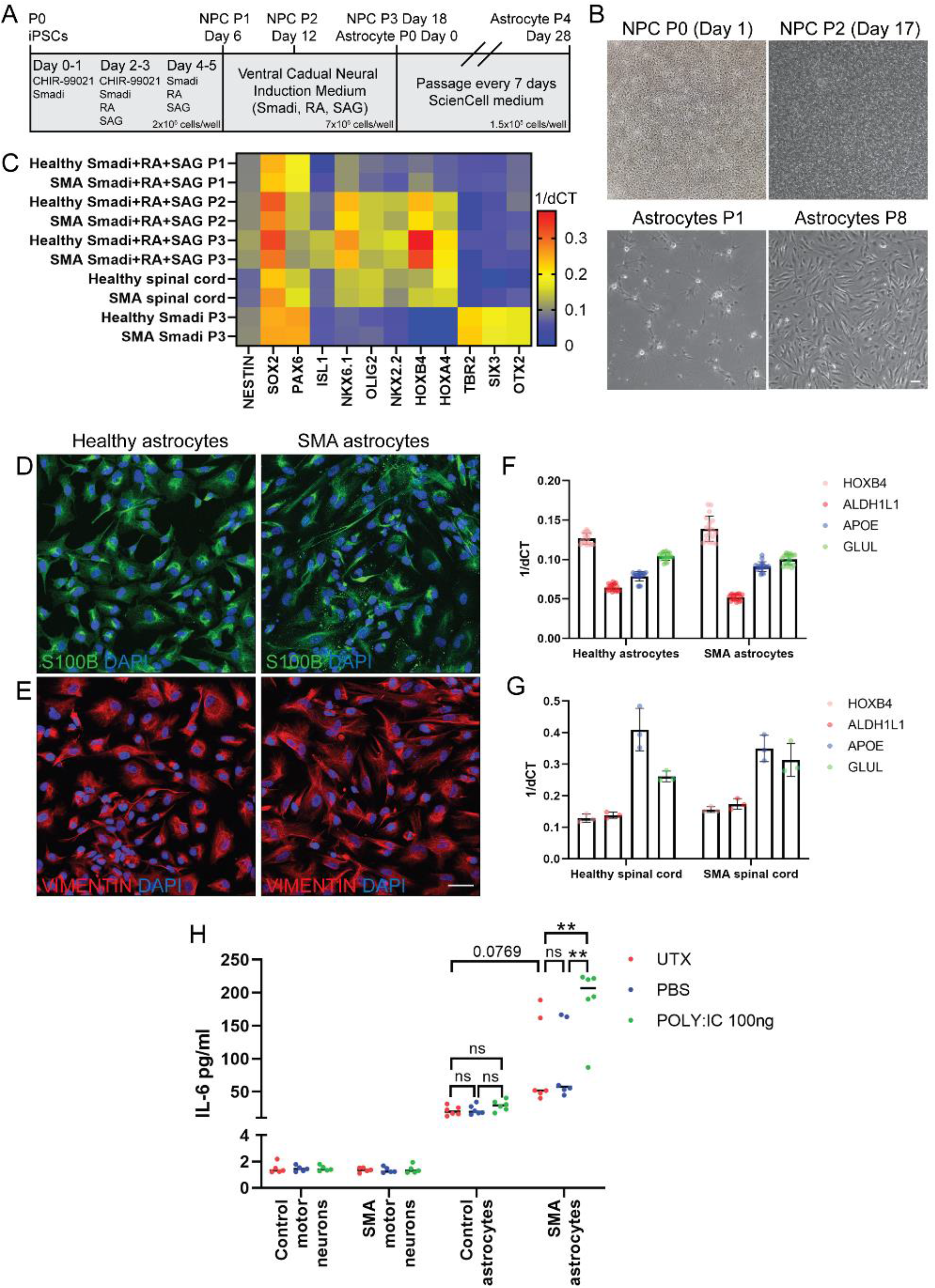
Spinal cord astrocyte differentiation of induced pluripotent stem cells (iPSCs). **(A)** Schematic of neural progenitor cell (NPC) and astrocyte differentiation protocols from iPSCs. NPCs are patterned and cultured for Day 18 before beginning astrocyte differentiation. Astrocytes beyond P4 (Day 28) are considered fully differentiated. **(B)** Representative brightfield images of NPC and astrocyte cultures during early and later stages of culture. Scale bar: 100μm. **(C)** Early addition of Wnt agonist (CHIR 99021), SMAD inhibition (SB431542 and LDN-193189), retinoic acid (RA) and hedgehog smoothened agonist (SAG) during NPC differentiation (P1-3) increases the expression of general NPC-associated genes (NESTIN, SOX2, PAX6) and spinal cord ventral caudal-associated genes (ISL1, NKX6.1, OLIG2, HOXB4 and HOXA4). Post-mortem spinal cord and ventral-caudal patterned NPC (P2 onwards) samples show comparable gene expression profiles, whereas Smadi only treated NPCs show increased forebrain transcription factor gene expression (TBR2, SIX3, OTX2). Markers characteristic of astrocyte identity, including S100B **(D)** VIMENTIN **(E)**, ALDH1L1, APOE, GLUL **(F)**, and spinal cord ventral caudal transcription factor HOXB4 (D) are readily detected in astrocytes differentiated from NPC cultures and in post-mortem spinal cord samples **(G)**. Scale bar: 50μm. **(H)** Significantly elevated levels of interleukin-6 (IL-6) are detected in conditioned media from stimulated (POLY:IC treated) SMA astrocytes, compared to untreated (UTX) and vehicle treated (PBS) SMA astrocytes, and equivalent healthy astrocyte samples. iPSC-derived motor neurons served as a negative control for this assay. 2-way ANOVA, Tukey’s multiple comparisons test. **pvalue<0.005.

SMA and Ctrl astrocytes were next differentiated from the ventral-caudal NPCs using astrocyte specific medium. A small neuronal population was noted during early stages of astrocyte differentiation (Figure 1B; Astrocyte P1), which diminished at later stages of culture (Figure 1B; Astrocytes P8). Ventral-caudal patterned astrocytes were found to express characteristic markers of astrocyte identity and function such as S100β (Figure 1D), VIMENTIN (Figure 1E), ALDH1L1, APOE, and GLUL (Figure 1F), which showed similar expression trends to human spinal cord samples (Figure 1G). iPSC-derived astrocytes and human spinal cord samples also showed equivalent expression of the spinal cord ventral-caudal transcription factor HOXB4 (Figure 1F, G). The Ctrl and SMA astrocytes were found to secrete interleukin 6 (IL6) into the media, with SMA astrocytes secreting slightly higher levels of IL6 at baseline (p=0.0769, ns, Figure 1H). When stimulated with polyinosinic-polycytidylic acid (poly:IC), a compound that activates inflammatory responses via toll-like receptors,^45^ Ctrl astrocytes slightly increased IL6 secretion though not significantly, whereas SMA astrocytes significantly upregulated IL6 secretion compared to untreated and PBS-treated SMA astrocyte samples (p<0.005, Figure 1H). These data fit with the findings from our previous model^30,32^ and establish these spinal cord astrocytes as phenotypically and functionally consistent with other *in vitro* models.^41,42,46,47^

### Differences in SMA astrocyte neurotrophic support and reactivity compared to healthy controls

Our lab previously reported decreased GDNF production and increased production of GATA6 in SMA astrocytes compared to healthy controls.^32^ These findings are consistent with our updated differentiation in which RNA transcripts of GDNF and BDNF were both significantly downregulated in the SMA astrocytes compared to Ctrl (Figure 2A, p=0.0033, 0.0221). GATA6 and NFκB transcripts were found to be significantly upregulated in the SMA cultures, also confirming our previous findings (Figure 2B, p<0.0001, p=0.0001). These transcript changes were maintained in protein expression as confirmed by immunocytochemistry (ICC) (p=0.0187, Figure 2C-E).

**Figure 2.**
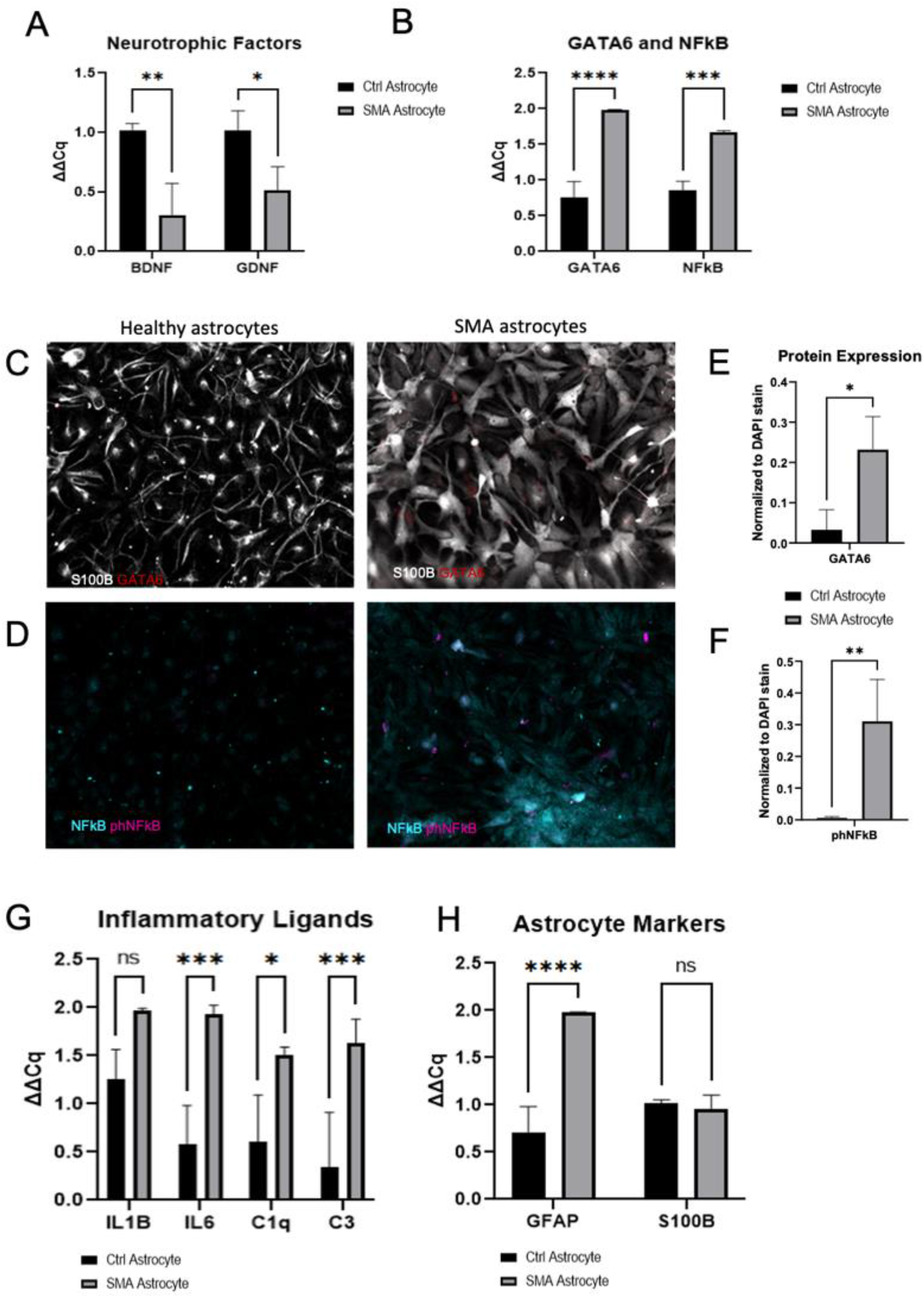
GATA6 and nuclear NFκB upregulation in SMA astrocytes. **(A)** Decreased neurotrophic support through BDNF, GDNF and **(B)** increased GATA6, NFκB transcript production confirmed in SMA astrocytes compared to healthy controls. **(C)** Representative images of GATA6 and **(D)** phNFκB protein expression in SMA and Ctrl Astrocytes by immunocytochemistry (**E, F** quantification). **(G)** Upregulation of transcripts for pro-inflammatory cytokines and complement factors (IL1β, IL6, C1q, C3) and **(H)** activation marker GFAP in SMA astrocytes compared to Ctrl, with no changes in general astrocyte marker S100β expression. 1-way ANOVA, Tukey’s multiple comparisons test. *pvalue<0.05, **<0.005, ***<0.0005, ****<0.0001.

Importantly, phosphorylated NFκB (phNFκB) was also found to be upregulated in the nuclei of SMA astrocytes compared to Ctrl (Figure 2D-F, IHC p=0.0031). phNFκB is known to enter the nucleus and regulate the gene expression of pro-inflammatory cytokines.^48,49^ Accordingly, we found upregulated transcripts for interleukin 1 beta (IL1β) and interleukin 6 (IL6), as well as upregulated complement cascade members C1q and C3 in the SMA astrocyte cultures (Figure 2G, p=0.0741, 0.0006, 0.0198, 0.0009). Glial fibrillary acidic protein (GFAP), a canonical marker of astrocyte activation, was also found upregulated, though none of these changes were associated with any changes in the general astrocyte marker S100B expression (Figure 2H, p<0.0001, 0.8529). Overall, these findings confirm the increased activation of SMA astrocytes compared to healthy control astrocytes in our updated culture system. They also identify GATA6 and NFκB as potential drivers of increased inflammatory ligand production in SMA.

### Manipulation of GATA6 associated with changes in reactivity and neurotrophic support

To directly examine the influence of GATA6 upregulation on driving astrocyte activation, we infected healthy control astrocytes (Ctrl) with a lentivirus expressing GATA6 (GATA6OE) or a GFP control. We also separately infected SMA astrocytes with a lentivirus expressing an shRNA against GATA6 (GATA6KD) construct or GFP control. We first confirmed GATA6 viral expression or knockdown in RNA transcripts by qRT-PCR (Figure 3A) and confirmed its protein expression by ICC (OE fold change of 9.7576, KD fold change of −3.2181, Figure 3C, E). Ctrl GATA6OE astrocytes showed increased GATA6 levels equivalent to SMA astrocyte levels (Ctrl GATA6OE to SMA GFP qPCR: p=0.9665, ICC p=0.2320), and SMA GATA6KD astrocytes were found to have decreased GATA6 levels similar to Ctrl astrocyte baseline (SMA GATA6KD to Ctrl GFP qPCR:0.0630, ICC p=0.9783). Interestingly, no significant changes in NFκB transcript levels were observed in Ctrl GATA6OE samples compared to Ctrl GFP samples (Figure 3B), although nuclear phNFκB protein levels were found significantly increased (ICC p=0.0178, Figure 3F). GATA6KD in the SMA astrocytes did have an associated significant decrease in NFκB transcript levels (p=0.0459) as well as a significant decrease in nuclear phNFκB (ICC p=0.0289, Figure 3B, 3D, 3F).

**Figure 3.**
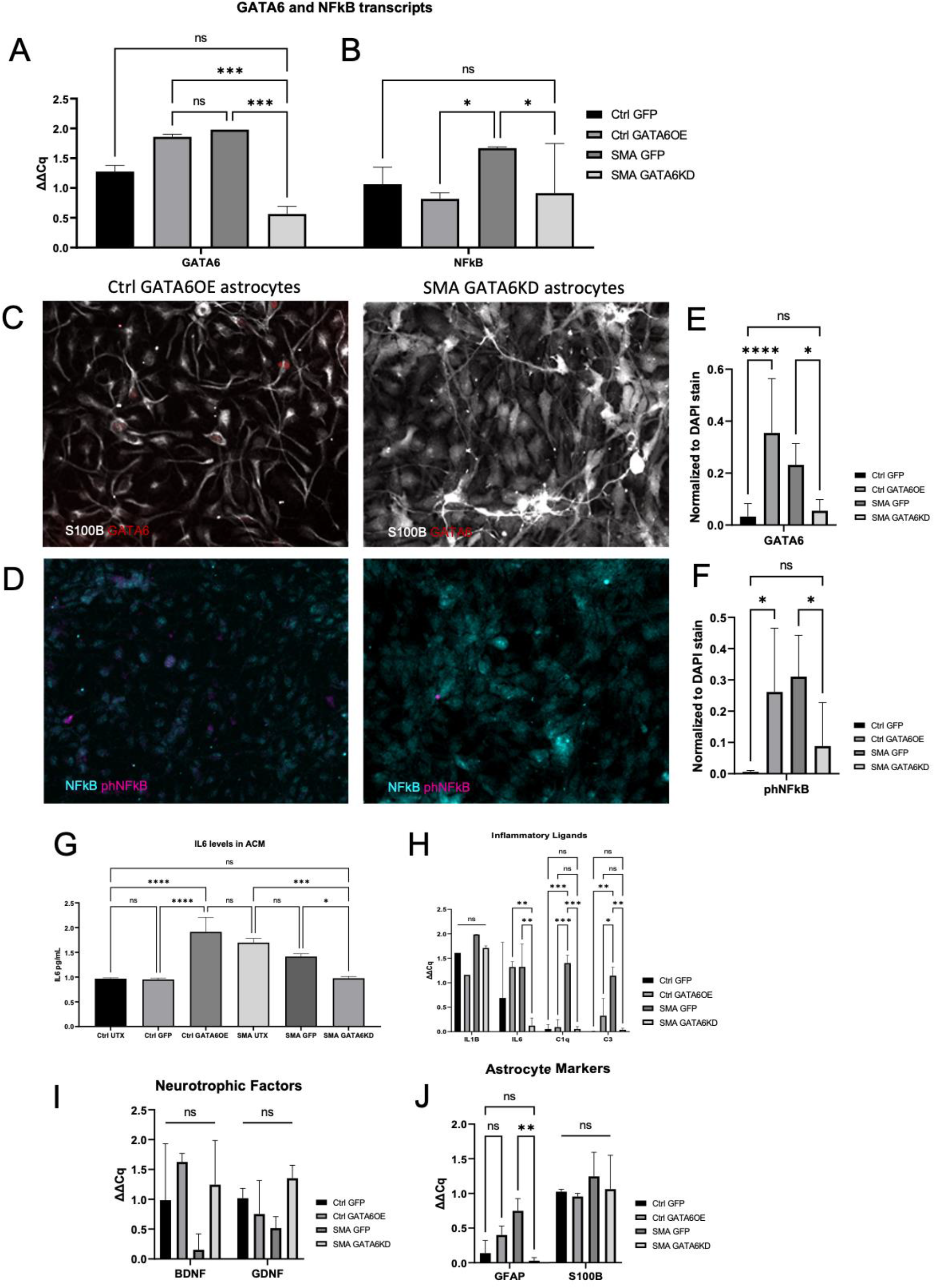
Changes in inflammatory and neurotrophic gene expression after astrocyte GATA6 manipulation. **(A)** Confirmation of GATA6 transcript reduction (SMA) or overexpression (Ctrl) after exposure to GFP control, GATA6 expressing, or GATA6KD lentivirus and **(B)** associated change in NFκB transcript production. **(C, D)** Confirmation of GATA6 and phNFκB protein expression changes after lentiviral manipulation assessed by ICC (**E, F** quantification). **(G)** Secreted IL6 protein levels confirm transcript changes after GATA6 manipulation. **(H)** Significant decrease in IL6, C1q, and C3 transcripts after GATA6KD in SMA astrocytes, trends for increases after GATA6OE in Ctrl astrocytes. **(I)** No significant changes in neurotrophic support after GATA6 manipulation. **(J)** Significant decrease in reactive GFAP transcripts, but importantly not in characteristic marker S100β. 2-way ANOVA, Tukey’s multiple comparisons test for qPCR; 1-way ANOVA, Tukey’s multiple comparisons test for ICC, *pvalue<0.05, **<0.005, ***<0.0005, ****<0.0001.

In relation to the NFκB data, we examined canonical pro-inflammatory cytokines in astrocyte samples, finding a significant increase in secreted IL6 after GATA6OE (Ctrl GFP to Ctrl GATA6OE p<0.0001, Figure 3G) and a significant decrease after GATA6KD (SMA GFP to SMA GATA6KD p=0.0002, Figure 3G). No significant changes in IL1B were detected (Figure 3H). Significant decreases in C1q (p=0.0004) and C3 (p=0.0035) transcript levels were also found after GATA6KD, and there was only a slight increase in expression after GATA6OE (fold change = 0.4286, 0.9898, ns) (Figure 3H). No significant changes in neurotrophic factors or astrocyte marker S100B were found in any condition (Figure 3I, 3J). GFAP transcript was significantly reduced after GATA6KD in SMA astrocytes (p=0.0087) but was not significantly increased in GATA6OE Ctrl astrocytes (fold change =0.6500, ns, Figure 3J). These findings suggest that aberrant GATA6 expression in SMA astrocytes is sufficient to modulate inflammatory ligands, complement components, and GFAP transcripts.

### Effect of astrocyte GATA6 manipulation on spinal motor neuron survival

We next investigated whether GATA6-associated reactivity and inflammation in astrocytes contributed to spinal motor neuron loss *in vitro*. Following the established Maury et al. protocol^50^ for generating spinal motor neurons (MNs) from human iPSCs, we matured plated MNs for 21-28 days before analysis. To examine the possible contribution of GATA6 to SMA astrocyte neurotoxicity, conditioned media (ACM) from Ctrl, Ctrl-GFP, Ctrl-GATA6OE astrocytes, as well as SMA, SMA-GFP, and SMA-GATA6KD astrocytes was applied onto Ctrl and SMA MNs. Cell death was then measured using a TUNEL assay and normalized to a live cell counterstain (7AAD, Figure 4A&B, D&E). Quantification of percent change was calculated using normalized values compared to one Ctrl UTX ACM replicate for Ctrl and SMA MNs (Figure 4C, F).

**Figure 4.**
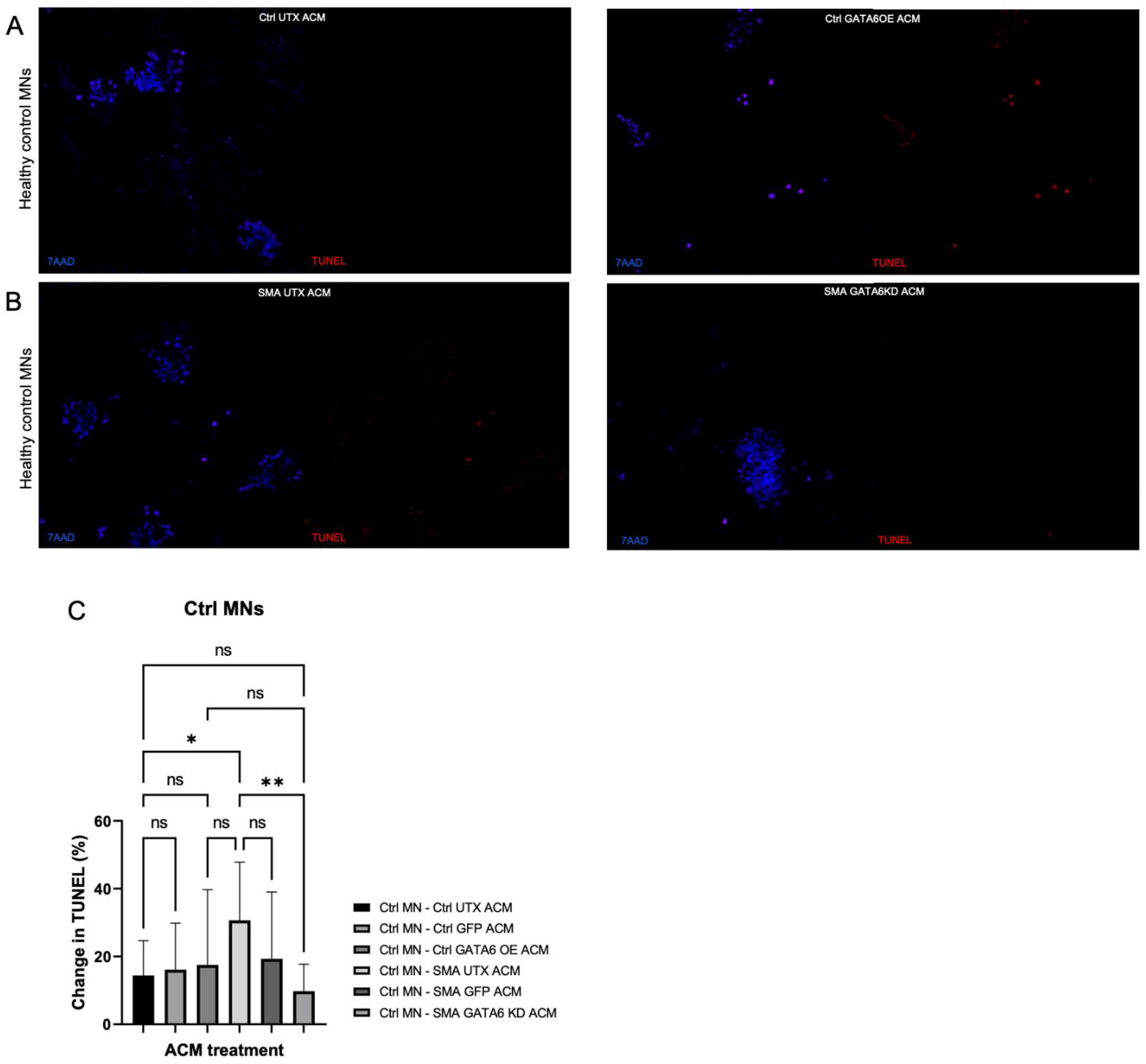
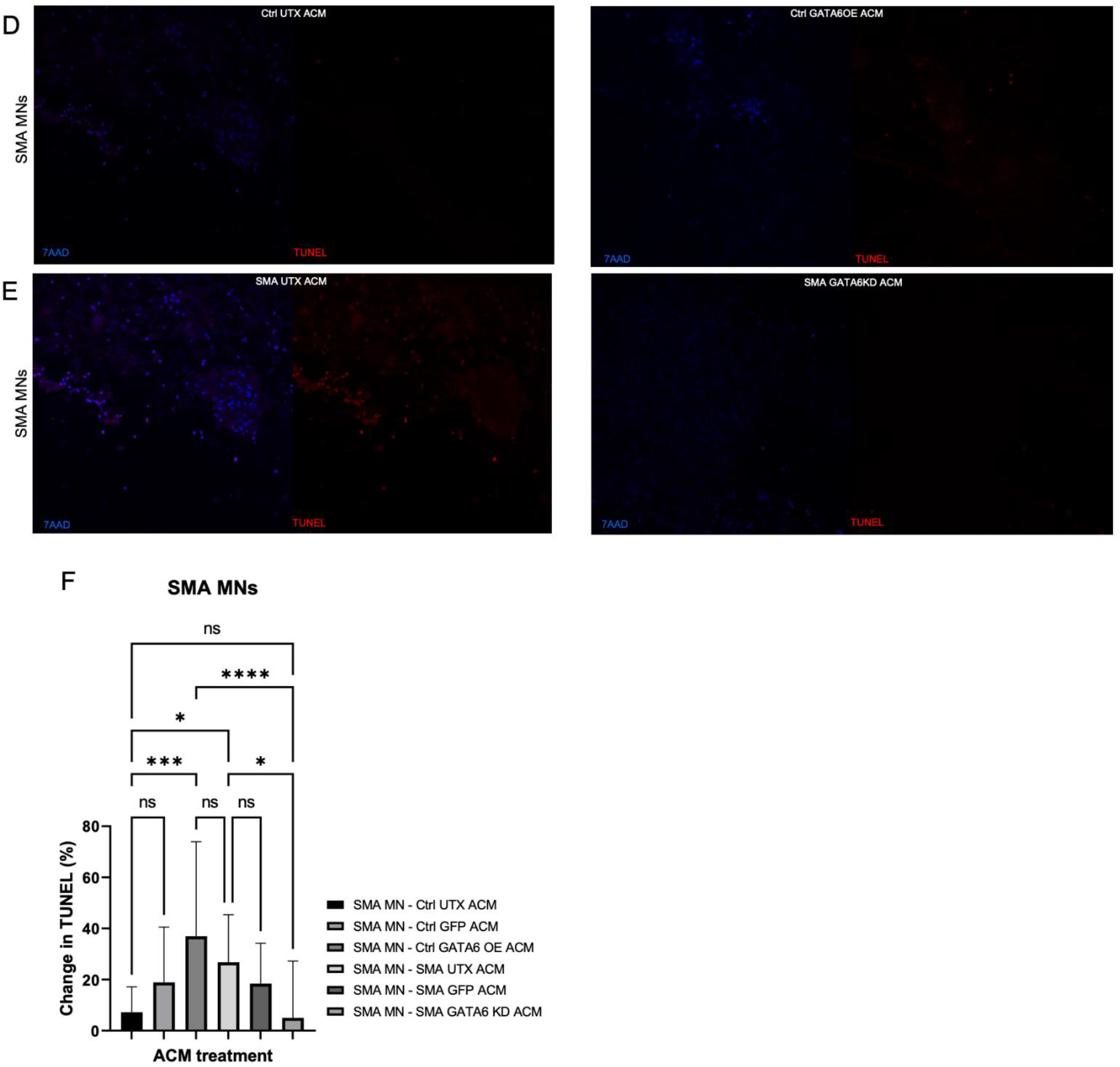
Spinal motor neuron survival after astrocyte GATA6 manipulation. Conditioned media from Ctrl untreated (UTX) or GATA6OE astrocytes applied onto Ctrl **(A)** and SMA **(D)** spinal motor neurons (MN). Media from SMA UTX or GATA6KD astrocytes also applied onto Ctrl **(B)** and SMA **(E)** MNs. Cell death was assessed with a TUNEL kit. TUNEL stain was normalized to live cell nuclear 7AAD stain and represented as percent change from the first WT UTX ACM replicate for Ctrl and SMA MNs **(C, F).** In Ctrl and SMA MNs, SMA UTX was significantly more toxic than Ctrl ACM, an effect which GATA6KD ameliorated. GATA6OE was significantly more toxic than Ctrl UTX only when applied onto SMA MNs. No differences in cell death were found between UTX and GFP conditions. 1-way ANOVA, Tukey’s multiple comparisons test. *pvalue<0.05, **<0.005, ***<0.0005, ****<0.0001.

Consistent with our previous findings, in both Ctrl and SMA MNs, ACM from untreated SMA astrocytes induced more cell death than ACM from untreated Ctrl (Ctrl UTX) astrocytes (p=0.0181, 0.0354, Figure 4A-4C, 4D-4F). No significant differences were found between the untreated and GFP-control samples in either Ctrl or SMA astrocytes (Ctrl-Ctrl UTX:GFP p=0.9992, Ctrl-SMA UTX:GFP 0.2805, SMA-Ctrl UTX:GFP p=0.4729, SMA-SMA UTX:GFP p=0.7920). GATA6OE ACM did significantly increase cell death compared to Ctrl UTX ACM in SMA MNs (p=0.0001), though not in Ctrl MNs (p=0.9850). This difference is likely due to the inherent vulnerability of SMA MNs. Strikingly, SMA GATA6KD ACM significantly decreased MN cell death in both Ctrl and SMA MNs compared to SMA UTX ACM (p= 0.0021, 0.0128). These data support the idea that GATA6 overexpression in SMA astrocytes drives neurotoxicity.

### Effect of astrocyte activation and GATA6 manipulation on microglial activation

Finally, we examined the effect of SMA astrocyte activation on human iPSC-derived microglia from SMA patients and healthy controls. Many models of neurodegeneration establish a role for activated microglia in driving further neuron loss.^27,51–56^ Because many of the increased pro-inflammatory cytokines and complement components found within our SMA astrocyte cultures are key signals for microglial activation and phagocytosis, we aimed to examine the possibility of SMA astrocyte-driven microglial activation contributing to spinal motor neuron loss. We started by generating microglia from SMA and Ctrl iPSC lines using established differentiation protocols and commercial kits (Figure 5A).^57,58^ Generated microglia were confirmed TMEM119+, P2RY12+, and Iba1+ before analyses (Figure 5B).

**Figure 5.**
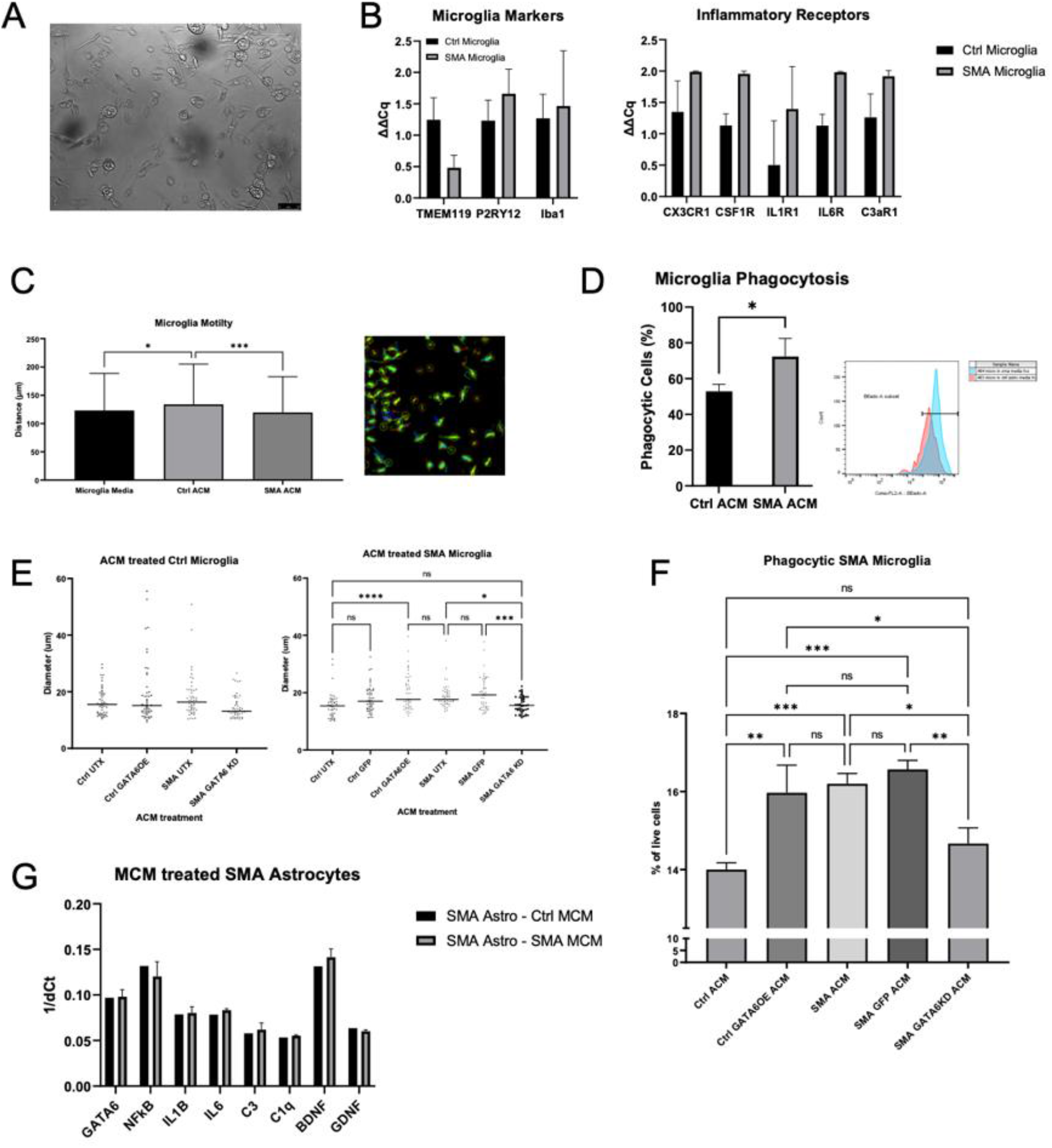
Microglial activation after exposure to astrocyte conditioned media (ACM). iPSC-derived microglia from Ctrl and SMA lines express microglia-specific markers TMEM119, P2RY12, and Iba1 **(A** representative image, **B)**. **(B)** SMA Microglia show trends for upregulated transcripts for inflammatory receptors including CX3CR1, CSF1R, IL1R1, IL6R, and C3aR1 (ns). **(C)** SMA microglia exposed to Ctrl ACM show increased motility compared to SMA ACM in live image tracking (left - representative image, right - quantification). **(D)** SMA Microglia show increased phagocytosis of FluoSpheres after SMA ACM treatment compared to Ctrl ACM. **(E**, left) No changes in phenotype (soma diameter) after Ctrl Microglia treated with GATA6 manipulated ACM. **(E**, right) A significant increase in soma size of SMA Microglia treated with GATA6OE compared to Ctrl UTX was found, as well as a significant decrease after GATA6KD compared to SMA UTX. **(F)** Significantly decreased phagocytosis of FluoSpheres when SMA Microglia treated with GATA6LD ACM compared to SMA UTX and SMA GFP ACM. **(G)** No changes found in any transcripts after SMA Astrocytes treated with microglia conditioned media. 1-way ANOVA, Tukey’s multiple comparisons test. *pvalue<0.05, **<0.005, ***<0.0005, ****<0.0001.

We first examined differences in inflammatory receptors at baseline between the Ctrl and SMA microglia (Figure 5B). Though no differences were significant, we did find trends for increased pro-inflammatory receptor transcripts including C3aR1, CXCR1, CSFRA, IL1R1, and IL6R in SMA microglia (Figure 5B). To test if SMA microglia may have heightened sensitivity to astrocyte secreted factors, we first applied Ctrl UTX and SMA UTX ACM onto SMA microglia generated from a yolk-sac lineage protocol (Figure 5C, 5D). Ctrl ACM increased microglia mobility—a sign of non-activated, surveilling microglia^18,59^ – compared to control media treated cells as measured by distance traveled through a live-image tracking assay (p=0.0316, Figure 5C; image). The application of SMA ACM onto SMA microglia significantly reduced this surveilling phenotype (p=0.0009, Figure 5C). To more directly examine functional differences, we measured phagocytic ability using fluorescent bead-based flow cytometry (Figure 5D). We found a significantly increased number of phagocytic SMA microglia after treatment with SMA ACM than Ctrl ACM (p=0.0350, Figure 5D).

We then wanted to examine the role of GATA6 in the SMA astrocyte-driven microglial activation phenotype and confirm our findings in a hematopoietic lineage model of microglia (Figure 5E, 5F). First, we found no differences in soma diameter – a sign of activation^60^ – between Ctrl UTX, Ctrl GATA6OE, or SMA UTX ACM applied onto Ctrl microglia (p=0.3193, 0.6850, ns, Figure 5E). A slight but significant decrease in diameter was seen in the SMA GATA6KD treated Ctrl microglia compared to Ctrl GATA6OE (p=0.0154, Figure 5E). In the SMA microglia treatments, a significant increase in size was seen in the Ctrl GATA6OE condition, comparable to SMA UTX treated microglia (Ctrl UTX:GATA6OE p<0.0001, GATA6OE:SMA UTX p=0.7106, Figure 5E). SMA microglia treated with SMA GATA6KD ACM showed a significant decrease in diameter, reducing sizes to Ctrl UTX levels (SMA UTX:GATA6KD p=0.0422, Ctrl UTX: GATA6KD p=0.9986). No differences were found between Ctrl UTX and Ctrl GFP, or SMA UTX and SMA GFP conditions (p=0.1295, 0.8410). Functionally, a significant decrease in phagocytic cells was found in SMA microglia treated with the SMA GATA6KD ACM compared to SMA UTX or SMA GFP ACM (p=0.0102, 0.0019, Figure 5F). A significant increase in phagocytic cells was also found in after treatment with the Ctrl GATA6OE ACM compared to Ctrl UTX ACM (p= 0.0014, Figure 5F).

Finally, we aimed to test if the activating or toxic effect was unique to SMA astrocytes or if SMA microglia had similar activating effects. Therefore, we treated SMA and control astrocytes with SMA or control microglia conditioned medium (MCM). Interestingly, no differences in activation were found when astrocytes were treated with media conditioned by Ctrl or SMA microglia (ns, Figure 5G). GATA6 transcripts were not differentially expressed between SMA and Ctrl microglia or between SMA and Ctrl motor neurons analyzed by RNAseq (data not shown) suggesting aberrant GATA6 expression is more astrocyte restricted. Overall, these data suggest a role for SMA astrocytes in SMA microglial activation and supports a role of aberrant GATA6 expression as a driving factor in astrocyte mediated neurotoxicity and microglial phagocytosis.

## Discussion

The loss of SMN in SMA causes selective lower motor neuron loss in patients, and this is replicated *in vivo* with SMA mouse models.^61^ However, in purified motor neuron cultures *in vitro*, SMA iPSC-derived motor neurons do not show increased cell death compared to healthy controls.^32,62^ This recent distinction, along with a body of evidence showing that targeting SMN restoration to only motor neurons has limited effect, suggest a role for non-cell autonomous contributions to SMA motor neuron vulnerability.^6,7,9,63^ In parallel, pre-synaptic deficits and aberrant activation of glial cells in SMA have been observed before overt motor neuron loss.^30^ Specifically, SMA astrocytes have been shown to decrease neurotrophic and growth factor production and increase the pro-inflammatory production of GFAP, IL6, and IL1B.^30,32,34^ The application of SMA astrocyte conditioned media has been shown to have detrimental effects on both healthy control MNs and those with reduced SMN.^32,64^ Activated (Iba1+, M1) microglia have also been found to be increased at early symptomatic stages and localized to motor neurons in SMA rodents.^37,65,66^ Synapse loss has been associated with microglial phagocytosis, with complement factors C1q and C3 implicated in targeting microglia removal of healthy synapses.^38,66^ Together, these data robustly indicate a role for glia in contributing to motor neuron loss in SMA.

Here, we utilized human iPSC-derived astrocytes, microglia, and spinal motor neurons from healthy control and SMA patient lines to examine disease properties in human cells. Though all astrocytes are thought to share common features (e.g., upregulation of GFAP after activation) and functions (e.g., formation and maintenance of the blood-brain barrier), recent studies have identified differences in gene expression, morphology, and function between astrocytes in distinct regions of the CNS.^39,67^ Even within the spinal cord, three distinct populations of astrocytes are hypothesized to preferentially support neurons from the same progenitor subtypes.^40,68,69^ To address these emerging ideas, we started by developing a more disease-relevant astrocyte culture by patterning neural progenitor cells with small molecules to mimic spinal cord development, then differentiated these into functional spinal cord astrocytes. ^39,40,70^ We found that SMA astrocytes upregulated classical markers of activation, decreased neurotrophic support, and increased pro-inflammatory ligands and complement components compared to controls. These data establish our spinal cord astrocytes as a disease-relevant and functional model concurrent with others being used in the field.^41,42,46,47^ They also allow us to examine astrocyte activation and interactions with spinal motor neurons and microglia in an SMA-specific context throughout the rest of this study.

In order to more specifically investigate the proposed relationship between SMN, GATA6, and NFκB in the context of astrocyte activation, we infected astrocytes with a lentivirus to drive GATA6 expression in Ctrl astrocytes or knock it down in SMA astrocytes. Interestingly, we found that though the transcript and protein levels of GATA6 were successfully manipulated, only the knockdown of GATA6 in SMA astrocytes, not the overexpression in Ctrl astrocytes, was associated with a significant change in NFκB transcript levels. Both, however, were associated with increased nuclear phNFκB protein levels. Knocking down GATA6 in the SMA astrocytes significantly decreased inflammatory and activation marker transcripts, though it did not restore neurotrophic support through BDNF or GDNF. Within these SMN-deficient SMA astrocytes, GATA6 does appear to be a main contributor of aberrant activation, confirming what transcriptome analyses, *in vivo* studies, and our own previous findings predicted.^32,36^ In contrast, overexpressing GATA6 in the Ctrl astrocytes showed trending—but not significant—increases in these transcripts. This suggests that these inflammatory ligands are more tightly regulated by NFκB than GATA6, with GATA6 potentially competing with functional alternative regulators of NFκB in healthy SMN-expressing astrocytes.^71^ The relationship between GDNF, BDNF, and GFAP may be similarly complicated. Histone deacetylase (HDAC), an enzyme which can increase DNA susceptibility to transcription factor (TF) regulation, has shown inverse associations with GFAP compared to BDNF and GDNF. In different contexts, low HDAC (DNA more accessible for TF regulation) is associated with decreased GFAP, whereas high HDAC (DNA less accessible for regulation) is associated with increased BDNF and GDNF.^72,73^ These complicated associations with the disease-specific genetic changes may begin to illuminate why the impact of GATA6KD was more striking in the SMA astrocytes than its overexpression in controls.

The effect of the GATA6 manipulated astrocytes on SMA and control MN survival follows a similar pattern. Expectedly, SMA ACM was found to increase motor neuron cell death in SMA and Ctrl cultures compared to Ctrl ACM, and the knockdown of GATA6 in SMA astrocytes ameliorated this effect in both SMA and Ctrl MNs. Strikingly, the overexpression of GATA6 in Ctrl astrocytes increased cell death only in the SMA MNs, not in the healthy controls. This again could be due to regulatory mechanisms—like those involving SMN—which are intact in the Ctrl MNs but are absent in SMA MNs leaving them more vulnerable to even slight changes in astrocyte secreted factors. One example of this is the SMN-deficiency-linked activation of p53 found in vulnerable motor neurons at pre-symptomatic stages in SMA mice.^32^ This distinction between GATA6OE toxicity to Ctrl and SMA MNs emphasizes the importance of therapeutically targeting the early-activating SMA astrocytes, especially in patients not receiving the AAV therapy to restore SMN. It also provides an interesting topic of future study regarding the resistance of AAV9-SMN treated motor neurons to astrocyte toxicity, specifically regarding GATA6-driven toxicity.

Next, we began to investigate the emerging idea of pathologically activated microglia in SMA by differentiating microglia from SMA and healthy control iPSCs. At baseline, we observed differences between the Ctrl and SMA microglia in pro-inflammatory receptors known to activate microglia from a neurotrophic, surveilling phenotype—characterized by small soma and highly motile processes—towards an ameboid, phagocytic phenotype.^23,24,59^ We also identified upregulated complement receptor C3aR1, a receptor which has been specifically tied to microglial activation and phagocytosis of neurons or synapses in several models of neurodegeneration.^27,74–76^ The upregulation of these receptors in SMA microglia may prime them for the upregulated ligands produced by SMA astrocytes to encourage activation. Encouragingly, we found that SMA ACM applied onto SMA microglia increased activated phenotypes compared to Ctrl ACM or no treatment in both yolk-sac lineage and hematopoietic progenitor lineage microglia. These data first confirmed the ability of iPSC-derived astrocytes and microglia to communicate via secreted factors. Additionally, they suggest that microglia-mediated phagocytosis could contribute to viable neuron loss in SMA, as has been suggested in neurodegenerative models such as Alzheimer’s and Parkinson’s disease.^77–79^ Importantly, the use of individually differentiated cultures and ACM treatments in this study suggest that microglia activation in SMA may be fueled by activated astrocyte secreted factors.

Finally, we examined the potential impact of GATA6 knockdown on this astrocyte-driven microglial activation. Morphologically and functionally, we found that treatment with SMA UTX and Ctrl GATA6OE ACM increased activated phenotypes of SMA microglia compared to Ctrl UTX or SMA GATA6KD ACM. The role of the complement proteins—especially C3—within this process is one promising area for future studies. C3 is upstream of complement cascade products which can be secreted to act as tags for phagocytosis (C3b, C5a), activators for immune cells (C3a), or initiators of the membrane attack complex (C5b).^80^ In SMA, already vulnerable MNs receiving neurotoxic conditioning and C5b signals from activated astrocytes may be upregulating their own C3 in addition to initiating the terminal pathway towards cell lysis.^81,82^ Excess complement products then may be additionally tagging nearby healthy neurons for phagocytosis-mediated cell death (i.e. phagoptosis) as well as functional synapses for removal by astrocyte-activated microglia. More experiments to examine the production of C3 by SMA MNs before and after ACM treatment, as well as in MN-microglia co-cultures, are needed to investigate this hypothesis. Nevertheless, these data demonstrate the ability of iPSC-derived astrocytes and microglia to communicate via secreted factors and provide compelling evidence for astrocyte-driven activation of microglia in SMA. Of particular interest, this activation appears to be an astrocyte-first mechanism, as the application of microglia conditioned media onto SMA astrocytes showed no effect This is in contrast to microglia-first activation theories which have been proposed in the context of Alzheimer’s, Huntington’s, and Parkinson’s disease.^83–85^ This is an area of future research, as C3-expressing activated astrocytes were observed in post-mortem tissue from patients with Alzheimer’s, Huntington’s, and Parkinson’s disease, as well as multiple sclerosis (MS) and amyotrophic lateral sclerosis (ALS).^86^ It would be especially of interest to compare the activation timelines of SMA and ALS, as motor neurons are particularly affected in these disorders— despite quite dissimilar genetic causes— and microglia have been directly linked to motor neuron loss in ALS.^87,88^ Using our isolated iPSC-derived culture system and treatments of ACM or MCM, we may be able to parse out if this astrocyte-first activation is specific to SMA, or if *in vivo* and post-mortem studies are missing early mechanisms of neuron loss.^89^

Together, these data provide evidence to support the predicted role of GATA6 in astrocyte activation and its neurotoxic effects on spinal motor neurons in SMA. They also suggest a role for astrocyte-driven microglia activation, which may contribute to MN death within SMA. More consideration for astrocytes and microglia when designing therapeutics for SMA may positively impact MN health to improve lifespan and function for patients.

## Materials and Methods

### Pluripotent Stem Cells

Two healthy control iPSC lines (21.8, 4.2) and three SMA patient iPSC (7.12, 3.6, 8.3) lines were utilized in these experiments.^32^ All pluripotent stem cells were maintained on Matrigel (Corning) in Essential 8 (Gibco) and passaged every 4-6 days. The iPSCs and differentiated cells were confirmed mycoplasma negative.

### Astrocyte differentiation

Spinal cord patterned astrocytes were generated from iPSC-derived neural progenitor cells (NPCs). Catalog numbers for each reagent are listed in supplemental materials (Table S1) and a protocol schematic is shown in Figure 1A. Briefly, iPSCs were grown to confluency, dissociated with Accutase, and plated at 2 million cells/well into Matrigel (Corning) coated 6 well-plates in NPC base medium (50% DMEM/ F12, 50% Neurobasal, 2% B27, 1% N2, 1% Antibiotic/ Antimycotic, 0.1% B-mercaptoethanol, 50ng/mL Laminin, and 0.5uM Ascorbic Acid) supplemented with 10uM Y-27632, 3uM Chir-99021, 40uM SB431542, and 0.2uM LDN193189. On day 1, NPC media was changed and supplemented with 3uM Chir-99021, 40uM SB431542, and 0.2uM LDN193189. Cells were washed with PBS if a lot of cell death was observed. Days 2 and 3 involved NPC media changes supplemented with 3uM Chir-99021, 40uM SB431542, 0.2uM LDN193189, 100nM retinoic acid (RA), and 500nM hedgehog smoothened agonist (SAG). From day 4 onwards, NPC media was supplemented with 40uM SB431542, 0.2uM LDN193189, 100nM RA, and 500nM SAG and changed daily. Cells were passaged via Accutase treatment on Day 6 (P1), Day 12 (P2), and Day 18 (P3). Plating density for NPC P1 and P2 is 7 million cells per well (of a 6 well plate) and plating density for NPC P3 to begin astrocyte differentiation (P0) is 150,000 cells/well (of a 6 well plate).

Astrocyte differentiations were cultured with ScienCell Astrocyte Medium containing 1% Astrocyte Growth Supplement, 1% Penicillin/Streptomycin, and 2% B27. Cells were fed every 48 hours and passaged with Accutase every 6-9 days upon confluency (minimum of 3 passages). Astrocyte P1, P2, and P3 were all plated at 150,000 cells/ well, at P4 cells were considered differentiated and split onto coverslips for IHC, into 6 well plates for RNA or protein, or into T75 tissue culture treated flasks for ACM generation and collection. Astrocyte conditioned media was collected upon each media change after P4, spun to remove any cells or debris, and stored in sterile Falcon tubes at –20C. Frozen medias were slowly thawed on ice before use. Poly:IC treatments were diluted to 100ng/mL in supplemented Astrocyte media and left for 24 hours. Cells were then rinsed with PBS and fed with supplemented Astrocyte media. Media was collected from stimulated astrocytes after 48 hours, and pellets were collected for analyses. IL-6 ELISA assay was performed on collected media (Affymetrix eBioscience, #88-7066-22).

### Motor neuron differentiation and treatments

Spinal motor neurons were differentiated based on the Maury et al. (2015) protocol.^50^ Briefly, embryoid bodies were generated from iPSCs and patterned in the presence of Chir-99021 with dual SMAD inhibition (SB 431542 and LDN 1931899) followed by treatment with retinoic acid (RA), smoothened agonist (SAG), and DAPT. Spinal motor neuron progenitor cells were then dissociated and plated on Matrigel-coated glass coverslips for terminal differentiation and maturation in growth factor supplemented medium for 28-42 days in vitro. Treatments with ACM were performed with 20% ACM in MN Maturation media after day 28 and left for 48 hours at 37C before fixing, collection, or analyses.

### Microglia differentiations

Microglia were differentiated using the protocols established by Haenseler et al. (2017)^57^ or the newly available differentiation kit by STEMCELL Technologies based on the McQuade et al. (2018) protocol^58^ (#05310, #100-0019, #100-0020). Briefly, iPSCs were patterned into macrophages (pMacs) using BMP4, VEGF, SCF, IL3, and M-CSF. pMacs were matured into microglia (pMGL) using N2, IL34, and GM-CSF (experiments shown in Figures 5C, 5D). Alternatively, iPSCs were differentiated into hematopoietic progenitor cells (HPCs) using the STEMdiff Hematopoietic Kit (StemCell Technologies). Floating HPCs were then collected and plated at 50,000 cells/mL in STEMdiff Microglia Differentiation media (StemCell Technologies) for 24 days, followed by rapid maturation in STEMdiff Microglia Maturation media (StemCell Technologies) for at least 4 days before usage (experiments shown in 5B, 5E, 5F, 5G). Microglia were confirmed TMEM119+, CD45+, and CD11b+ (Figure 5B).

### qRT-PCR

RNA was isolated from control and SMA cell pellets using the RNeasy Mini Kit (Qiagen) following manufacturer’s instructions, quantified using a Nanodrop Spectrophotometer, treated with RQ1 Rnase-free Dnase (Promega), and converted to cDNA using the Promega Reverse Transcription system (Promega). SYBR green qRT-PCR was performed in triplicate using cDNA and run on Bio-Rad CFX384 real time thermocycler. Primers for each target are available in Supplemental materials (Table S2). Cq values for each target gene were normalized to housekeeping gene (GAPDH) values and Ct values for each sample were normalized to a human cortex sample to calculate relative fold change (ddCT) in gene expression.

### Immunocytochemistry

Plated cells were fixed in 4% paraformaldehyde (PFA) for 20 minutes at room temperature and rinsed with PBS. Nonspecific labeling was blocked and the cells permeabilized with 0.25% Triton X-100 in PBS with 1% BSA and 0.1% Tween 20 for 15 minutes at room temperature. Cells were incubated with primary antibodies overnight at 4C, then labeled with appropriate fluorescently-tagged secondary antibodies. Hoechst nuclear dye was used to label nuclei. Primary antibodies used were chicken anti-Vimentin (abcam, ab24525), mouse anti-S100β (Sigma, S2532), rabbit anti-GATA6 (Cell Signaling #5851), mouse anti-NFκB p65 (Cell Signaling #6956), and rabbit anti-phospho-NFκB p65 (Cell Signaling #3033). Secondary antibodies used were donkey anti-mouse AF488 (Invitrogen), goat anti-chicken AF568 (Invitrogen), donkey anti-rabbit AF546 (Invitrogen), and donkey anti-mouse AF647 (Invitrogen). All primaries were used at a 1:500 dilution, and all secondaries were used at 1:1000 dilution. Analyses were performed on three randomly selected fields per coverslip using standard fluorescent microscopy using equivalent exposure conditions. Total fluorescence in each channel was measured using FIJI (ImageJ) software.

### TUNEL assay

Plated cells were fixed in 4% PFA for 20 minutes at room temperature, rinsed with PBS, and then stained using a TUNEL Assay Kit - BrdU-Red (abcam, ab66110). Manufacturer’s instructions were adapted for coverslips instead of Coplin jars. Briefly, cells were permeabilized as described for ICC, washed, and incubated with DNA labeling solution for 1 hour at 37C. Cells were washed and optional 7AAD nuclear counterstain was then applied for 30 minutes at room temperature. Coverslips were imaged within 3 hours of staining with standard fluorescent microscopy. Three images were acquired from randomly selected fields for each coverslip. Images were analyzed for total fluorescence in either channel using FIJI (ImageJ) software. Representative images were acquired on a Zeiss confocal microscope within 3 hours of staining.

### Microglia assays

Microglia were treated with 1:2 astrocyte conditioned media with microglia maturation media for 24 hours at 37C. Live image tracking was performed with 1:6000 Invitrogen Cell Tracker Green CMFDA dye with 5,000 cells per condition. Images were acquired over 16 hours using an Olympus IX83 Inverted Light Microscope and analyzed on ImageJ using OlympusViewer, TrackMate, and Chemotaxis Tool plugins. Soma size measures were generated by fixing cells onto glass coverslips and staining with Iba1 (abcam, ab5076), then acquiring three random images per coverslip. Images were blindly analyzed using Nikon Elements Advanced Research package to measure soma diameter of total n=50 cells per condition. Phagocytosis assay was performed by treating cells with astrocyte conditioned media or microglia maturation media (control) for 24 hours at 37C. Cells were then incubated with FluoSpheres Sulfate Microspheres (ThermoFisher #F8851) for 15 minutes at 37C. Unbound beads were removed from cell suspension, then cells and bound beads were suspended in FACS flow buffer on ice. Flow cytometry was performed immediately and gated on live cells to remove any unbound beads from downstream analyses. Flow cytometry was performed on an LSRFortessa X-20 Special Order Research Product or BD Accuri C6 (BD Biosciences, Franklin Lakes, NJ).

### Statistical analyses

Experimental conditions within each experiment were performed in technical triplicates for a minimum of three independent experiments. Data were analyzed using GraphPad Prism software and the appropriate statistical tests including the Student’s t-test, 1-way ANOVA, and 2-way ANOVA followed by Tukey’s post hoc analysis of significance. Changes were considered statistically significant when p<0.05.

## Acknowledgements

We would like to acknowledge the Children’s Research Institute FLOW Core at the Medical College of Wisconsin and thank Melissa Whyte for her help with the microglia flow cytometry assays. GATA6 knockdown and overexpression constructs were generously provided by Dr. Stephen Duncan, and lentiviruses were generated in the Blood Research Institute/ MCW Viral Vector Core. We thank Dr. Michele Battle for helpful GATA6 discussions. This work was supported by grants to A.D.E. from the National Institutes of Health (R21NS102911) and Cure SMA. B.G.B is supported by the NIH (R01NS119594).

## Author Contributions

R.L.A, E.W., and G. K. performed and analyzed experiments. R.L.A, E.W., A.D.E., G.K., and B.G.B designed experiments and interpreted data. A.D.E. and B.G.B supervised the study and provided funding. R.L.A wrote the manuscript, E.W. and R.L.A created figures and legends, and E.W. and A.D.E. edited the manuscript.

