## Supplemental tables 1 and 2 for "Viral mediated knockdown of GATA6 in SMA iPSC-derived astrocytes prevents motor neuron loss and microglial activation"

Table S1 – Spinal cord astrocyte differentiation materials

| Product | Company | Catalog Number |
|---|---|---|
| Matrigel | Corning | 356234 |
| DMEM | ThermoFisher | 11965092 |
| F12 | ThermoFisher | 11765054 |
| Neurobasal | ThermoFisher | 21103049 |
| B27 supplement | ThermoFisher | 17504044 |
| N2 supplement | ThermoFisher | 17502048 |
| Antibiotic/ Antimycotic | ThermoFisher | 15240062 |
| B-mercaptoethanol | ThermoFisher | 21985023 |
| Laminin | Millipore Sigma | L2020 |
| Ascorbic Acid | Millipore Sigma | A4544 |
| Y-27632 | Selleck Chemical | S1049 |
| Chir-99021 | Selleck Chemical | S1263 |
| SB 431542 | Selleck Chemical | S1067 |
| LDN 193189 | Selleck Chemical | S7507 |
| Retinoic Acid (RA) | Millipore Sigma | R2625 |
| Smoothened Agonist (SAG) | Millipore Sigma | 566660 |
| Acctuase | StemCell Technologies | 7922 |
| Astrocyte Medium | ScienCell | 1801 |
| polyinosinic-polycytidylic acid (poly:IC) | InvivoGen | tlrl-pic, tlrl-pic-5 |

Table S2 – Primers for qRT-PCR

| Cell Type | Target gene | Forward sequence | Reverse sequence |
|---|---|---|---|
| NPC | NESTIN | TCAAGATGTCCCTCAGCCTGGA | AAGCTGAGGGAAGTCTTGGAGC |
| NPC | Sox2 | GCTACAGCATGATGCAGGACCA | TCTGCGAGCTGGTCATGGAGTT |
| NPC | Pax6 | TGGTATTCTCTCCCCCTCCT | TAAGGATGTTGAACGGGCAG |
| NPC | ISL1 | GCAGAGTGACATAGATCAGCCT | GCCTCAATAGGACTGGCTACCA |
| NPC | NKX2.2 | CCTTCTACGACAGCAGCGACAA | ACTTGGAGCTTGAGTCCTGAG |
| NPC | HOXB4 | CTGGATGCGCAAAGTTCACGTG | CGTGTCAGGTAGCGGTTGTAGT |
| NPC | HAXA4 | GGCAAGGAGCCCGTGGTGTAC | TCCTTCTCCAGCTCCAAGACCT |
| NPC | TBR2 | AAATGGGTGACCTGTGGCAAAGC | CTCCTGTCTCATCCAGTGGGAA |
| NPC | SIX3 | ACCGGCCTCACTCCCACACA | CGCTCGGTCCAATGGCCTGG |
| NPC | OTX2 | ACAAGTGGCCAATTCACTCC | GAGGTGGACAAGGGATCTGA |
| Astrocyte | BDNF | CATCCGAGGACAAGGTGGCTTG | GCCGAACTTTCTGGTCCTCATC |
| Astrocyte | GDNF | CGCCGAAGACCGCTCCCTCG | ATCCATGACATCATCGAACTGATC |
| Astrocyte | GATA6 | GCCACTACCTGTGCAACGCCT | CAATCCAAGCCGCCGTGATGAA |
| Astrocyte | NFkB | GCAGCACTACTTCTTGACCACC | TCTGCTCCTGAGCATTGACGTC |
| Astrocyte | IL1B | CCACAGACCTTCCAGGAGAATG | GTGCAGTTCAGTGATCGTACAGG |
| Astrocyte | IL6 | AGACAGCCACTCACCTCTTCAG | TTCTGCCAGTGCCTCTTTGCTG |
| Astrocyte | C1q | CAACACAGGCTGCTACGGGATC | CTGCCCTTTGGGTCCTCGGAT |
| Astrocyte | C3 | GTGGAAATCCGAGCCGTTCTCT | GATGGTTACGGTCTGCTGGTGA |
| Astrocyte | GFAP | GTCCCCCACCTAGTTTGCAG | TAGTCGTTGGCTTCGTGCTT |
| Astrocyte | S100B | AACAAAGGAGGACCTGAGAGTAC | CTTTGCCATCTCTACACTGGTCC |
| Microglia | TMEM119 | GGATAGTGGACTTCTTCCGCCA | GGAAGGACGATGGGTAATAGGC |
| Microglia | P2RY12 | TGCCAAACTGGGAACAGGACCA | TGGTGGTCTTCTGGTAGCGATC |
| Microglia | Iba1 | CCCTCCAAACTGGAAGGCTTCA | CTTTAGCTCTAGGTGAGTCTTGG |
| Microglia | CX3CR1 | CACAAAGGAGCAGGCATGGAAG | CAGGTTCTCTGTAGACACAAGGC |
| Microglia | CSF1R | GCTGCCTTACAACGAGAAGTGG | CATCCTCCTTGCCCAGACCAAA |
| Microglia | IL1R1 | GTGCTTTGGTACAGGGATTCCTG | CACAGTCAGAGGTAGACCCTTC |
| Microglia | IL6R | GACTGTGCACTTGCTGGTGGAT | ACTTCCTCACCAAGAGCACAGC |
| Microglia | C3aR1 | CCTGCTGATGTGGTCTCACCTA | CCTTGTGGTAGCTCAGACTCGT |
